# Examining the potential involvement of NONO in TDP-43 proteinopathy in *Drosophila*

**DOI:** 10.1101/2023.12.13.571416

**Authors:** Rafael Koch, Emi Nagoshi

**Affiliations:** Department of Genetics and Evolution and Institute of Genetics and Genomics of Geneva (iGE3), University of Geneva, CH-1205 Geneva, Switzerland

**Keywords:** Nuclear bodies, Neurodegenerative disorders, Locomotor deficits, Lifespan

## Abstract

The misfolding and aggregation of TAR DNA binding protein-43 (TDP-43), leading to the formation of cytoplasmic inclusions, emerge as a key pathological feature in a spectrum of neurodegenerative diseases, including amyotrophic lateral sclerosis (ALS) and frontotemporal lobar dementia (FTLD). TDP-43 shuttles between the nucleus and cytoplasm but forms nuclear bodies (NBs) in response to stress. These NBs partially colocalize with nuclear speckles and paraspeckles that sequester RNAs and proteins, thereby regulating many cellular functions. The laboratory of Steven Brown has recently found that the non-POU domain-containing octamer-binding protein (NONO), a component of paraspeckles, forms novel nuclear speckle-like structures in mouse cortical neurons in response to stress and sleep deprivation. These findings suggest the possibility of a functional link between NONO and TDP-43, potentially contributing to TDP-43 proteinopathy. Here, we demonstrate that loss of function in the *Drosophila* homolog of *NONO, no on or off transient A* (*NonA)*, exacerbates pathological phenotypes caused by *TDP-43* gain of function, leading to locomotor defects and life span shortening. These results provide supporting evidence for the functional link between NONO and TDP-43 and lay the foundation for dissecting underlying mechanisms.

## Introduction

TAR DNA binding protein-43 (TDP-43) is a highly conserved protein belonging to the heterogeneous nuclear ribonucleoprotein (hnRNP) family. Similar to other hnRNPs, TDP-43 contributes to various processes in nucleic acid metabolism, such as RNA splicing, mRNA stability, microRNA maturation, and transcriptional regulation (Buratti & Baralle, 2012). The presence of cytoplasmic inclusions containing TDP-43 has been identified in the nervous tissues of patients affected by a broad range of neurodegenerative disorders, including amyotrophic lateral sclerosis (ALS), frontotemporal lobar dementia (FTLD), Alzheimer’s disease (AD) (Neumann *et al*., 2006). Growing evidence supports TDP-43 misfolding as a key pathogenic factor in these disorders, leading to their collective designation as TDP-43 proteinopathy (de Boer *et al*., 2021). Despite the large body of studies, however, the pathogenic mechanism underlying TDP-43 proteinopathy remains incompletely understood.

TDP-43 shuttles between the nucleus and cytoplasm, predominantly localized in the nucleus under non-stress conditions. Recent studies have shown that, in response to stress, TDP-43 forms nuclear bodies (NB), the membraneless nuclear structures concentrating nuclear proteins, RNAs, and RNPs. These TDP-43 NBs partially colocalize with nuclear speckles and paraspeckles (Dykstra & Barmada, 2023). Although the native function of nuclear bodies remains elusive, the dynamic formation of TDP-43 NB formation in response to stress suggests its stress-mitigating role. The formation of NBs can alleviate the generation of cytoplasmic inclusions, and dysfunction in this process may contribute to the development of neurodegenerative disorders (Wang *et al*., 2020).

Paraspeckles are formed by the interaction between the long non-coding RNA *NEAT1* (nuclear-enriched abundant transcript 1) and the members of the *Drosophila* Behavior Human Splicing (DBHS) family of proteins, namely P54NRB/ non-POU domain-containing octamer-binding protein (NONO), splicing factor proline-glutamine rich (PSPC1), and PSF/SFPQ (Fox *et al*., 2018). Paraspeckles sequester proteins and RNAs, thereby regulating many cellular functions and cellular differentiation (Wang & Chen, 2020). A component of paraspeckle NONO interacts with and modulates the activities of Period (PER) proteins, the transcriptional repressors constituting a negative feedback loop of the circadian clocks. Reduction in NONO levels therefore impairs circadian gene expression in mammalian cells (Brown *et al*., 2005). The *Drosophila* homolog of NONO, encoded by the *no on or off transient A* (*nonA*) has a parallel role in the *Drosophila* circadian clock, regulating circadian gene expression and locomotor behavior through interacting with the circadian transcriptional activator CLOCK (Brown *et al*., 2005; Luo *et al*., 2018). While NONO regulates the circadian clockwork, it also modulates the outputs of circadian clocks in various tissues in a context-dependent manner, such as circadian-gating of cell cycle and wound healing (Kowalska *et al*., 2013) and post-transcriptional regulation of feeding/fasting cycle-dependent gene expression in the liver (Benegiamo *et al*., 2018).

Interestingly, NONO forms speckle-like structures in the nuclei, which are likely distinct from paraspeckles, in the hepatocytes upon feeding (Benegiamo *et al*., 2018). Furthermore, the laboratory of Steven Brown found that NONO forms these novel nuclear speckles in mouse cortical neurons in response to excitotoxic stress or sleep deprivation and facilitates RNA processing (personal communication from Steven Brown). Considering the dynamic subcellular localization of TDP-43, these findings collectively raise an intriguing possibility that NONO and TDP-43 may functionally interact to contribute to TDP-43 proteinopathy. This study begins to address this question using *Drosophila*, the model organism in which the roles of NONO and TPD-43 have been successfully studied (Brown *et al*., 2005; Elden *et al*., 2010; Li *et al*., 2010; Diaper *et al*., 2013; Luo *et al*., 2018). We show that NONO loss of function exacerbates pathological phenotypes caused by the TDP-43 gain of function, namely lifespan reduction and locomotor deficits. These results lay the foundation to further explore the functional link between NONO and TDP-43.

## Materials and Methods

### Drosophila strains

Flies were kept on standard corn meal/molasses food at 25°C in a 12h/12h light-dark (LD) regime. *elav-GAL4* is described previously (Kozlov *et al*., 2020). The following stocks were obtained from the Bloomington *Drosophila* Stock Center: *UAS-hTDP-43* (RRID:BDSC_79587), *nonA* RNAi 1 (TRiP#HMC03675, RRID:BDSC_52933), *nonA* RNAi 2 (TRiP#HMJ23111, RRID:BDSC_61279), *UAS-nonA RNAi* TRiP# HMC04383 (RRID:BDSC_56944), *D42-GAL4* (RRID:BDSC_8816), *GMR-Gal4* (RRID:BDSC_1104), *GMR57C10-GAL4* (RRID:BDSC_39171), *tubulin-Gal4* (RRID:BDSC_5138), and *UAS-mCD8::GFP* (RRID:BDSC_32185). *UAS-nonA RNAi* GD26411 was obtained from the Vienna Drosophila Resource Center (VDRC).

### Climbing assay

Climbing assays were performed as described previously (Bou Dib *et al*., 2014) with minor modifications. Flies were grouped into ~20 individuals and assessed groupwise from day 1 to 21 of age, every 7 days. Dead flies were not replaced. On the day of the assay, flies were transferred into graduated 100 ml measuring cylinders divided into 5 equally spaced zones. After a 1-h recovery from mild CO_2_ anesthesia, flies were tested by tapping them down and then allowing them to climb for 20 seconds. Each assay was repeated twice with a 5-minute break between repeats. Climbing assays were always performed at 2 h after lights-on (ZT2) to avoid the influence of circadian behavioral rhythms.

### Lifespan assay

Male flies of the same genotype were kept at a density of approximately 20 per vial, containing standard corn meal/molasses food. The vials were maintained in a controlled ~ 80% humidity at 25°C and under 12h/12h LD cycles. The vials were positioned horizontally to prevent flies from sticking to the food. Flies were transferred to fresh vials every two to three days and dead flies were counted. Kaplan-Meyer survival probability curves were plotted using the GraphPad Prism software (Version 10).

### Statistics

Climbing data were analyzed using one-way ANOVA with Tukey’s Honest Significant Difference (HSD) test in R software. The Kaplan-Meyer estimator and survival analysis log-rank tests were performed using the Mantel-Cox approach in GraphPad Prism software (Version 10). For all experiments, the level of significance was set at p<0.05.

### Genotypes

To control for the UAS copy number, an inert *10xUAS-mCD8::GFP* transgene was added when only *UAS-nonA RNAi* or *UAS-hTDP-43* was driven with the GAL4.

For climbing assay (Figure 1): *w*^*1118*^; *UAS-hTDP-43/+. w*^*1118*^; *+; D42-GAL4. w*^*1118*^; *UAS-hTDP-43/+; UAS-mCD8::GFP/D42-GAL4. w*^*1118*^; *UAS-RNAi 1/+; UAS-mCD8::GFP/D42-GAL4. w*^*1118*^; *UAS-RNAi 2/+; UAS-mCD8::GFP/D42-GAL4. w*^*1118*^; *UAS-RNAi 1/UAS-hTDP-43; D42-GAL4/+*.

**Figure 1.**
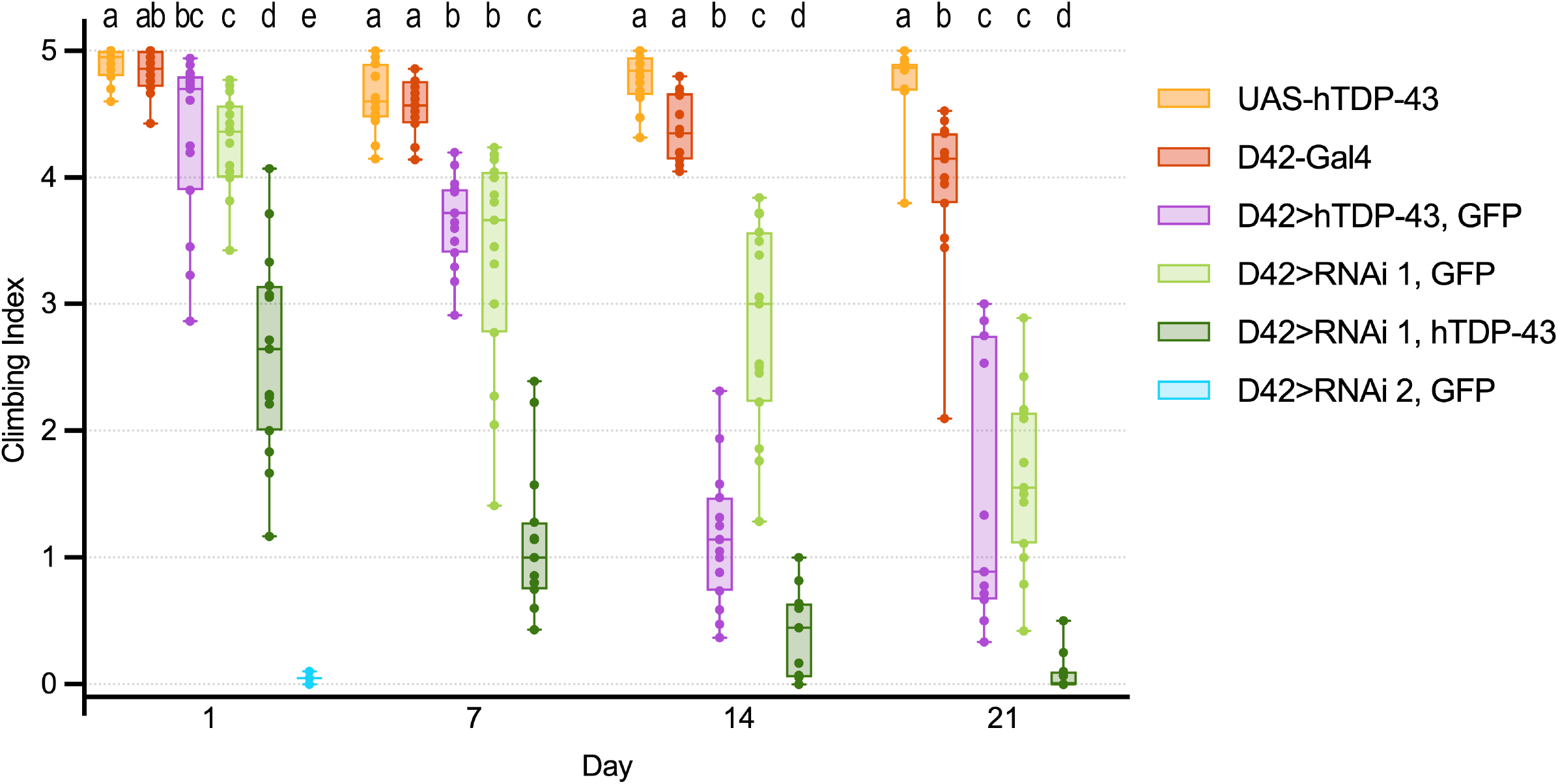
Knockdown of *nonA* in motor neurons exacerbates the locomotor deficits resulting from the overexpression of *hTDP-43*. Climbing ability was assessed in flies expressing *hTDP-43* or *nonA* RNAi from *D42-GAL4* independently or in combination, along with the control flies carrying only the GAL4 driver or *UAS-hTDP-43*, between age day 1 to 21. *UAS-mCD8::GFP* (GFP) was co-expressed when only one of the UAS transgenes was driven. Each dot represents the climbing ability of a group of ~20 males of indicated genotypes. N=5 independent experiments. The numbers of flies on day 1 were: *UAS-hTDP-43* (100), *D42>hTDP-43, GFP* (97), *D42>RNAi 1, GFP* (108), *D42>RNAi 1, hTDP-43* (74), and *D42>RNAi 2, GFP* (20). All three repeats per day are shown. *D42>RNAi 2* was semi-lethal and the limited number of individuals obtained could be tested on day 1 only (N=1). Different letters indicate significant differences between groups (p<0.05, one-way ANOVA with Tukey’s post-hoc test for multiple comparisons).

For lifespan analysis (Figure 2): *w*^*1118*^; *+; elav-GAL4/+. w*^*1118*^; *UAS-hTDP-43/+. w*^*1118*^; *UAS-RNAi 1/+. w*^*1118*^; *UAS-RNAi 2/+. w*^*1118*^; *UAS-hTDP-43/+; UAS-mCD8::GFP/elav-GAL4. w*^*1118*^; *UAS-RNAi 1/+; UAS-mCD8::GFP/elav-GAL4. w*^*1118*^; *UAS-RNAi 2/+; UAS-mCD8::GFP/elav-GAL4. w*^*1118*^; *UAS-RNAi 1/UAS-hTDP-43; elav-GAL4/+. w*^*1118*^; *UAS-RNAi 2/UAS-hTDP-43; elav-GAL4/+*.

**Figure 2.**
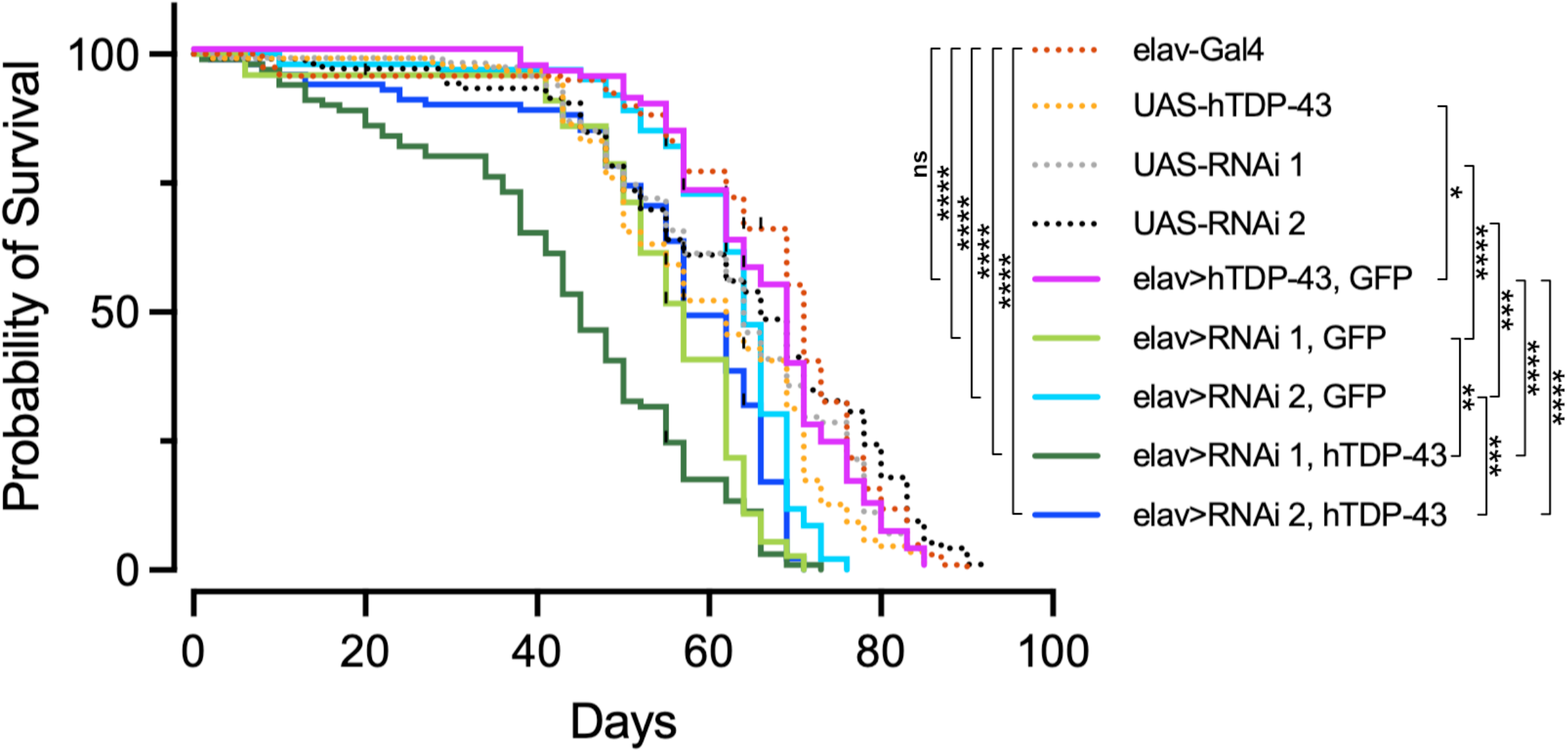
Overexpression of *hTDP-43* exacerbates the reduced lifespan resulting from *nonA* loss of function. *UAS-nonA RNAi* and *hTDP-43* were driven with a pan-neuronal *elav-GAL4*, individually or in combination. Lifespans of male flies of 5 to 6 independent replicates indicated genotypes were assessed at 25°C. The numbers of flies were: *elav-Gal4* (119), *UAS-hTDP-43* (125), *UAS-RNAi 1* (114), *UAS-RNAi 2* (106), *elav>hTDP-43, GFP* (95), *elav>RNAi 1, GFP* (49), *elav>RNAi 1, GFP* (101), *elav>RNAi1, hTDP-43* (101), and *elav>RNAi 2, hTDP-43* (102). Controls are plotted in dotted lines. ns, not significant. \**p*<0.05, \*\**p*<0.01, \*\*\**p*<0.001 and \*\*\*\**p*<0.0001 (Kaplan-Meier estimator).

For eye morphology analysis (Figure 3): *w*^*1118*^; *UAS-hTDP-43/GMR-GAL4; UAS-mCD8::GFP/+. w*^*1118*^; *UAS-RNAi 1/GMR-GAL4; UAS-mCD8::GFP/+. w*^*1118*^; *UAS-RNAi 2/GMR-GAL4; UAS-mCD8::GFP/+. w*^*1118*^; *UAS-RNAi 1, UAS-hTDP-43/GMR-GAL4; UAS-mCD8::GFP/+*.

**Figure 3.**
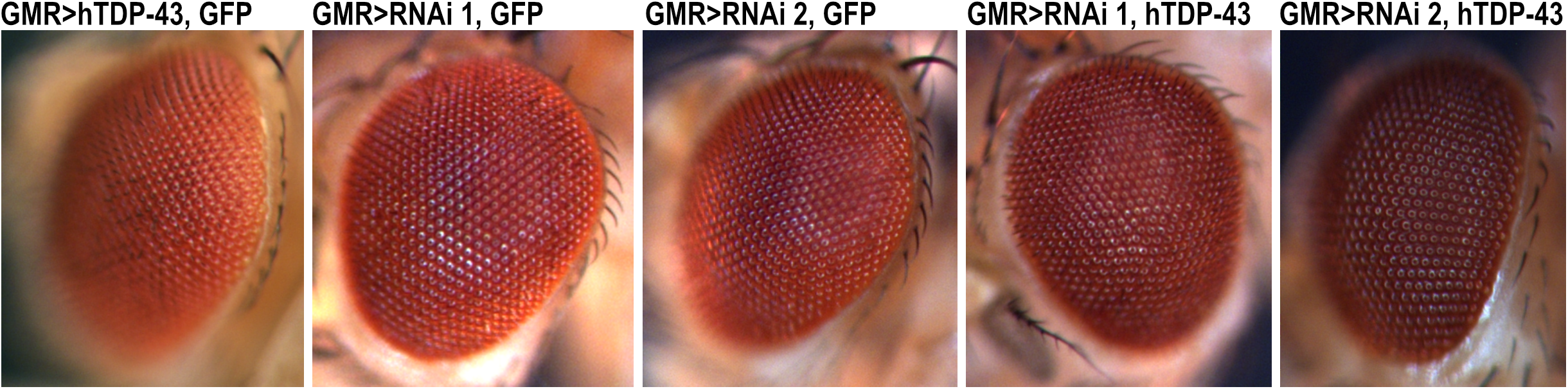
Expression of *nonA* RNAi or *hTDP-43* did not induce discernible eye defects. *UAS-nonA RNAi* and *hTDP-43*, either alone or in combination, were driven by *GMR-GAL4*. Eye morphology of 7-day-old adults was examined, revealing no ommatidial or necrotic defects. Representative images are shown.

## Results

Previous studies have demonstrated the pathophysiological effects of TDP-43 gain of function on cellular and organismal levels by the overexpression of human TDP-43 (hTDP-43) in flies (Elden *et al*., 2010; Li *et al*., 2010). To test if NONO influences the pathogenesis of TDP-43 proteinopathy, we wanted to combine hTDP-43 overexpression and NONO loss of function. To this end, we first assessed four independent UAS-RNAi lines (TRiP HMC03675, TRiP HMJ23111, TRiP HMC04383, and VDRC GD26411) for their efficiency in silencing endogenous *nonA* expression.

Ubiquitous expression of TRiP HMC03675 (referred to as RNAi 1) or TRiP HMJ23111 (referred to as RNAi 2) with *tubulin-GAL4* resulted in embryonic lethality, mirroring the phenotype of the *nonA* null allele (Stanewsky *et al*., 1993). In contrast, expressing VDRC GD26411 or TRiP HMC04383 with *tubulin-GAL4* did not affect viability. Driving *nonA* RNAi 1 with a pan-neuronal GMR57C10 driver was viable, whereas *nonA* RNAi 2 was mostly late pharate lethal. Expressing *nonA* RNAi 2 with a weaker pan-neuronal driver, *elav-GAL4* (Kozlov *et al*., 2020) was viable. These results indicate that *nonA* RNAi 1 and 2 have higher knockdown efficiency than the VDRC GD26411 and TRiP HMC04383 lines, with RNAi 2 exhibiting slightly higher efficacy than RNAi 1. Therefore, we chose to use *nonA* RNAi 1 and 2 in subsequent experiments.

The expression of *hTDP-43* in motor neurons was reported to induce progressive locomotor dysfunction (Elden *et al*., 2010; Li *et al*., 2010). We next tested if *nonA* RNAi modulates this phenotype by expressing *hTDP-43* together with *nonA* RNAi in motor neurons and assessing locomotion using the negative geotaxis (climbing) assays across ages. To control for the UAS copy number, an inert *UAS-mCD8::GFP* transgene was added when only *UAS-nonA RNAi* or *UAS-hTDP-43* was driven with the GAL4. Control flies carrying the motor neuron-specific *D42-GAL4* (Sanyal, 2009) only or UAS-*hTDP-43* only exhibited high levels of climbing ability at least up to age 21 days. Consistent with the previous studies, flies expressing *hTDP-43* with *D42-GAL4* displayed a significant progressive loss of climbing ability. Some *hTDP-43* expressing flies were unable to eclose from their pupal case and many larvae pupariated at the bottom in the food, reminiscent of the reported reduced larval motility (Li *et al*., 2010). Flies expressing *nonA* RNAi 1 in motor neurons were viable during the experimental period and displayed a progressive locomotor decline as compared to controls carrying only the GAL4 driver or one copy of the *UAS-hTDP-43* transgene. Flies expressing *nonA* RNAi 2 *D42-GAL4* were weak and showed strong locomotor impairments; the few eclosed individuals were unable to fly or climb and died before reaching day 7 (Figure 1).

Co-expressing *hTDP-43* and *nonA* RNAi 1 severely exacerbated locomotor deficits of flies expressing *hTDP-43* alone from day 1 and continuously up to at least day 21. Climbing assays of flies co-expressing *hTDP-43* and *nonA* RNAi 2 in motor neurons were not possible, as they were pharate lethal or were unable to eclose from the pupal case (Figure 1).

Pan-neuronal knockdown of *nonA* either with RNAi 1 or RNAi 2 under the control of *elav-GAL4* led to a reduced life span. Curiously, flies with RNAi 2 lived longer than those expressing RNAi 1, the inverse of the phenotype severity observed when they were expressed with *D42-GAL4*. While a previous study reported the lifespan reduction when *hTDP-43* was driven with *elav-GAL4* (Elden *et al*., 2010), we observed no differences in lifespan in *elav>hTDP-43* flies compared to the GAL4-only or UAS-only controls. Co-expression of *nonA* RNAi and *hTDP-43* further reduced life span compared to the expression of *nonA* RNAi alone. (Figure 2).

The expression of hTDP-43 in the eye with the *GMR-GAL4* driver was reported to cause progressive ommatidia loss, which was visible by day 6 (Elden *et al*., 2010; Li *et al*., 2010). In the same genotype, *GMR-GAL4* driving *hTDP-43*, we did not observe any eye degeneration phenotypes at day 7. Additionally, the expression of *nonA* RNAi 1 or 2 individually or with *hTDP-43* in the eye did not visibly affect the morphology of the eye at least by day 7 (Figure 3).

## Discussion

The pathophysiology of TDP-43 proteinopathy encompasses a broad spectrum, including motor and cognitive dysfunction, as well as a reduction in lifespan (Kawakami *et al*., 2019; de Boer *et al*., 2021). Here we investigated the potential functional interaction between NONO, a multifunctional RNA-binding protein, and TDP-43, with implications for the pathophysiology of TDP-43 in *Drosophila*. We found that loss of *nonA*, the fly homolog of *NONO*, exacerbates locomotor deficits induced by targeted expression of hTDP-43 in motor neurons. Furthermore, the lifespan shortening induced by the pan-neuronal loss of *nonA* is aggravated when co-expressed with hTDP-43. Taken together, our results provide support for the hypothesis that TDP-43 and NONO functionally interact within the nervous system, contributing to the modulation of TDP-43 pathology.

While the toxicity associated with hTDP-43 expression was recapitulated in our climbing assays, we did not observe the previously reported retinal degeneration by hTDP-43 overexpression driven by *GMR-GAL4* (Elden *et al*., 2010; Li *et al*., 2010). A potential explanation for this discrepancy could be that our flies were assessed at a constant 25°C and not shifted to 29°C after eclosion, a method employed in a previous study to enhance GAL4 activity (Elden *et al*., 2010). Additionally, we co-expressed an inert *UAS-mCD8::GFP* with *hTDP-43* to control for UAS copy number differences between *hTDP-43* flies and those co-expressing *NonA* RNAi. This may have caused a dilution effect that led to lower hTDP-43 expression levels. Notably, Elden et al. showed that a YFP fusion of hTDP-43 that has higher steady-state expression led to more severe eye defects. Therefore, it appears that the eye degeneration phenotype is sensitive to hTDP-43 dosage.

Similarly, the reduced life span was observed in *elav>hTDP-43* flies in the study by Elden et al., but not in our study. This discrepancy is also likely attributed to differences in experimental conditions, with flies kept at 29°C in Elden et al. and at 25°C in our study. In summary, the effect of genetic manipulation of NONO and TDP-43 on neuropathology is sensitive to the choice of the driver and experimental conditions.

Our findings lay the foundation for further investigation into the functional interaction between TDP-43 and NONO. Future studies examining their dynamic subcellular localization in both single and combined mutant genotypes, along with exploring their synergistic effects on neuronal morphology and function, may yield valuable insights.

## Acknowledgments

We are grateful to Steven Brown for sharing us with unpublished data and contributing to the conceptualization of this project. This study was supported by the grant from the Swiss National Science Foundation to E.N. (310030_189169).

## Author contribution

E.N. and R.K. conceived the project and planned the experiments. R.K. conducted the experiments and analyzed the data. R.K. and E.N. wrote the manuscript.

## Data Accessibility

All data in this study are displayed in the manuscript.

